# pH dependent inhibition from ammonium ions in the Pseudomonas mevalonii HMG-CoA Reductase crystallization environment

**DOI:** 10.1101/2020.06.03.132290

**Authors:** Vatsal Purohit, C. Nicklaus Steussy, Anthony R. Rosales, Chandra J. Critchelow, Tim Schmidt, Paul Helquist, Olaf Wiest, Andrew Mesecar, Cynthia V. Stauffacher

**Author notes:** AUTHOR INFORMATION Corresponding Author Department of Biology, Purdue University, Hockmeyer Hall of Structural Biology, 240 S. Martin Jischke Dr., West Lafayette, IN 47907.

## Abstract

HMG-CoA reductase (*Pseudomonas mevalonii*) utilizes mevalonate, coenzyme A (CoA) and the cofactor NAD in a complex mechanism involving two hydride transfers with cofactor exchange, accompanied by large conformational changes by a 50 residue subdomain, to generate HMG-CoA. Details about this mechanism such as the conformational changes that allow intermediate formation, cofactor exchange and product release remain unknown. The formation of the proposed intermediates has also not been observed in structural studies with natural substrates. Having been shown to be an essential enzyme for the survival of gram-positive antibiotic resistant pathogenic bacteria, studying its mechanism in detail will be beneficial in developing novel antibacterials. The enzyme has been shown to be catalytically active inside the crystal with dithio-HMG-CoA and NADH but curiously is found to be inactive in the reverse direction in the structure bound to mevalonate, CoA and NAD.

To understand the factors limiting activity in the HMGR crystal with mevalonate, CoA and NAD, we studied the effect of crystallization components and pH on enzymatic activity. We observed a strong inhibition in the crystallization buffer and an increase in activity with increasing pH. We attribute this inhibitive effect to the presence of ammonium ions present in the crystal since inhibition is also observed with several other ammonium salt buffers. Additionally, the lack of inhibition was observed in the absence of ammonium. The effect of each ligand (mevalonate, CoA and NAD) on the rate of the enzymatic reaction in the crystallization environment was further investigated by measuring their K_m_ in the crystallization buffer. The K_m_ measurements indicate that the hydride transfer step between NAD and mevalonate is inhibited in the crystallization environment. To test this further, we solved a crystal structure of pmHMGR bound to the post-hydride transfer intermediate (mevaldehyde) and cofactor Coenzyme A. The resulting turnover with the formation of a thiohemiacetal indicated that the crystallization environment inhibited the oxidative acylation of mevalonate and the reaction intermediate mevaldyl-CoA.

## Introduction

Class II HMG-CoA reductase *from P. mevalonii* (pmHMGR) catalyzes the oxidative acylation of mevalonate to form HMG-CoA, which is further converted into acetoacetate^[1-3]^. Since *P. mevalonii* utilizes R-mevalonate as a carbon source, the oxidative acylation of mevalonate is the predominantly occurring reaction in pmHMGR, unlike other homologues of this enzyme that have been characterized in other prokaryotic and eukaryotic organisms that favor the reduction of HMG-CoA for isoprenoid biosynthesis^[4-6]^. The enzyme undergoes two hydride transfer steps at the active site using 2 NAD^+^ cofactors, which results in the formation of two proposed intermediates, mevaldyl-CoA and mevaldehyde (Scheme 1).

**Scheme 1.**
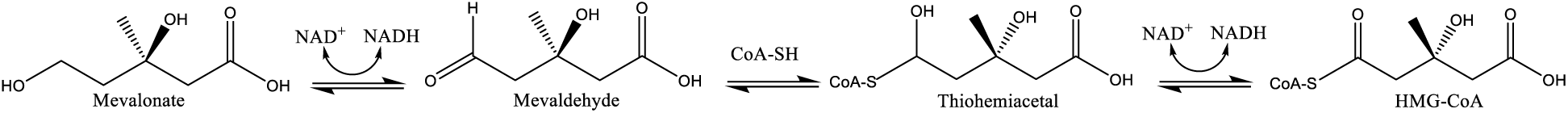
Proposed HMGR reaction mechanism in *P. mevalonii*

Crystallographic studies of pmHMGR have shown that the enzyme exists as a homodimer (90,000 g / mol) with the catalytic site formed at the dimer interface^[7, 8]^. Each monomer has a domain region that binds to cofactors coenzyme A (Large domain) and NAD (Small domain) with the substrate mevalonate binding at the homodimer interface^[9]^.

Movement of the 50-residue C-terminal region of the enzyme, the flap domain, has been found to be essential in the formation of the enzyme’s active site. The flap domain has been shown to form close contacts with the ligand-bound region upon substrate and cofactor binding to position the catalytic histidine (HIS 381) for catalysis^[8]^.

While biochemical and computational studies have hinted at the possibility of a bound mevaldehyde intermediate, it has not been observed in structural studies so far^[10, 11]^. Structures obtained with a slow substrate dithio-HMG-CoA have indicated the formation of mevaldyl-CoA in the crystal^[9]^. We also have structural information about the following stages of the reaction mechanism shown in Scheme 2.

**Scheme 2.**
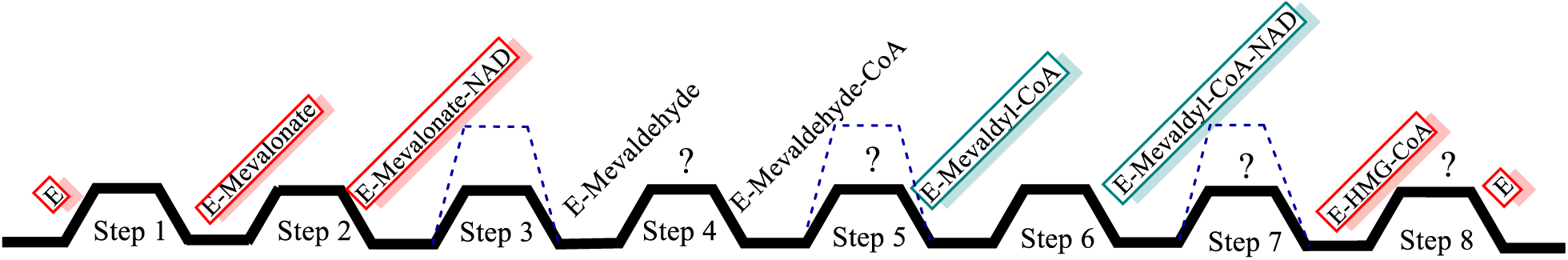
Reaction coordinate diagram showing the different steps within the HMG-CoA reductase catalytic reaction. Structures obtained with substrates, cofactors and products are shown in red and the ones that show evidence for intermediates with slow-substrates are shown in green. The dashed blue lines represent transition states during hydride or proton transfer steps along the reaction pathway.

Crystallographic structures of the enzyme bound to the substrate, mevalonate, and product, HMG-CoA have been observed. Additionally, the presence of the thiohemiacetal intermediate, mevaldyl-CoA has also been shown using crystallographic structures of the enzyme bound to a slow substrate, dithio-HMG-CoA^[9]^. However, we do not have structural evidence to show the formation of the other intermediate, mevaldehyde, the changes and structural arrangements that allow mevaldehyde and mevaldyl-CoA formation, the binding of the second NAD cofactor and the release of the product HMG-CoA.

The mevalonate pathway has been shown to be critical for the survival of antibiotic resistant gram-positive pathogens^[5, 12, 13].^ HMGR in *Pseudomonas mevalonii* has been shown to have a highly conserved active site fully comparable to its homologues in two pathogens that have shown antibiotic resistance, *Staphylococcus Aureus* and *Enterococcus Facaelis*^*[14]*^. Understanding the catalytic mechanism and the structural changes that facilitate it in more detail can be helpful in developing potent inhibitors that can be used as novel antibacterials. Studies using the slow substrate thio-HMG-CoA have shown that the enzyme is able to undergo turnover inside the crystal^[9]^, allowing the observation of trapped intermediates along the reaction pathway.

However, when trying to observe the turnover in the crystal with mevalonate, CoA and NAD in this study, we find that while the structures of the enzyme with these ligands show large conformational changes with the closure of the C-terminal region (flap domain) that allow for the formation of the catalytic active site, they do not indicate any product or intermediate formation.

To understand the cause for the lack of turnover in the quaternary complex within the pmHMGR crystal, where we would expect the reaction to go from mevalonate to HMG-CoA, we have asked the following questions in this study. (i) Which crystal constituents (if any) affect pmHMGR activity in this direction? (ii) Does the crystallization pH also play a role in the lack of turnover observed in crystal structures of pmHMGR bound to mevalonate, Coenzyme A and NAD? (iii) What ligand-enzyme interactions are being affected in crystals obtained with the quaternary complex?

## EXPERIMENTAL PROCEDURES

### pH profile measurements

The maximum turnover of the enzyme was measured at different pH values. The cofactors NAD^+^ and Coenzyme A for this assay were purchased from Sigma Aldrich. To prepare mevalonate, we purchased mevalanolactone from Sigma Aldrich and incubated the aqueous solution at pH 2 by treating it with HCl to break the lactone ring and stored it in aqueous solution at pH 7. To measure the turnover of the enzyme, the reaction for conversion of mevalonate to HMG-CoA was run in solution in various buffer conditions from pH 4-11. Substrates and cofactors in the final reaction were mevalonate at 4 mM, CoA at 0.51 mM and NAD^+^ at 2 mM. The substrates mevalonate, CoA and NAD were added to a 96-well plate with an assay volume of 100 µL. Substrates and cofactors in the final reaction were mevalonate at 4 mM, CoA at 0.51 mM and NAD^+^ at 2 mM. 100 µg of HMG-CoA reductase (*P. mevalonii*) which was expressed and purified using a previously described method was added to the well before the activity was measured ^[15-17]^. The absorbance of the samples was measured using a BioTek Synergy H1 Hybrid reader. The concentration of NADH produced was calculated from the absorbance values using the following equation: *conc* 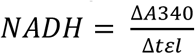 where ΔA340 denotes the linear increase in the absorbance after enzyme-ligand mixing. Δt is the time for which the absorbance was measured, ε is the extinction coefficient of NADH (6022 M ^−1^cm^−1^) and l (0.3 cm) is the path length of the microplate reader well. The reduction of NAD to NADH is observed via the measurement of emission peak from the nicotinamide moiety at 340 nm. The turnover of the enzyme was then measured using the following equation: 

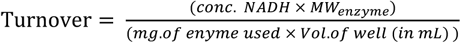

Here the molecular weight of enzyme, which exists as an obligate dimer is 90,000 g/mol^[7]^.

This measurement identifies the production of the reduced cofactor (NADH) in the two reaction steps that produce mevaldehyde from mevalonate and HMG-CoA from mevaldyl-CoA using a hydride transfer mechanism from the substrate mevalonate to the cofactor NAD and from the intermediate mevaldyl-CoA to another NAD molecule. While calculating the rate of change of absorbance indicates the formation of mevaldehyde, it cannot, with any certainty indicate the formation of the thioester bond in mevaldyl-CoA and its subsequent conversion to HMG-CoA.

The enzyme’s turnover at various pH values was plotted for different buffer conditions. The activity was measured at pH values from 4-10 in the enzymatic assay buffer (100 mM Tris, 100 mM Potassium Phosphate and 100 mM Glycine), crystallization buffer (1.2 M Ammonium sulfate, 100 mM ADA and 32% glycerol), Ammonium Chloride buffer (1.2 M Ammonium Chloride, 100 mM ADA and 32 % glycerol) and Potassium Sulfate buffer (0.27 M Potassium Sulfate, 100 mM ADA, 29 % glycerol).

### K_m_ measurements in crystallization conditions

To determine the K_m_ of the substrate and cofactor involved in the reaction converting mevalonate into HMG-CoA within the pmHMGR crystal environment, the rate in µmol / min / mg was measured at different concentrations for NAD, Coenzyme A and mevalonate in the crystallization buffer with pmHMGR. The concentration ranges used for CoA, NAD and mevalonate were 0-0.25 mM, 0-2.0 mM and 0-2.0 mM respectively. For the K_m_ measurements for each ligand, we kept the concentration of the other ligands used in the reaction at a saturated concentration of 0.51 mM (Coenzyme A), 2 mM (NAD) and 4 mM (mevalonate). The data was plotted and fit using a Michaelis Menten and Lineweaver Burk plot using the Sigma Plot 14.0 software to calculate the Km of each ligand.

### Diffraction measurements to study substrate / intermediate interactions

### pmHMGR bound to mevalonate, Coenzyme A and NAD

Purified HMGR (at 20 mg/mL) was crystallized using a sitting drop vapor phase exchange method. Crystal trays were set up using a reservoir solution with 1.2 M ammonium sulfate, 100 mM sodium ADA and 10 % glycerol at pH 6.7. A 200 x 250 µm crystal was used to prepare a seeding solution of crushed reductase crystals. 0.9 µL of the seeding solution that was diluted 10^3^-10^4^ fold and was added to 10 µL of reservoir solution, 10 µL of purified HMGR and 2.5 µL of nanopure water. To cryoprotect these crystals, the concentration of glycerol was increased to 30 %. 100 uL of 1 mM CoA, 1 mM NAD and 5 mM mevalonate were slowly introduced to the crystals by gradually replacing the reservoir solution and the substrates were allowed to soak in for at least 4 hours in order for them to bind strongly enough to observe them clearly in the crystallographic density. Crystallographic data from crystals with dimensions ≥ 250 x 250 x 100 µm extended to 2.2 Å. Diffraction data with these crystals were obtained at the Advanced Photon Source, Argonne National Laboratory (23-ID-B).

### pmHMGR bound to mevaldehyde and Coenzyme A

Mevaldehyde was prepared using a mevaldic acid precursor (the hemi-[N,N’-dibenzylethylenediammonium] salt of mevaldic acid) purchased from Sigma Aldrich^[18]^. HMGR crystals of size 400 x 500 x 500 µm in 1.2 M (NH_4_)_2_SO_4,_ ADA and Glycerol at pH 6.7 were transferred by capillary into another sitting drop well with the same solution. Mevaldehyde and CoAwere gradually introduced into the solution to obtain a final ligand concentration of 4 mM for both after exchanging out the starting solution. The crystals were soaked in ligand solution for 14 hours followed by cryoprotection by gradually introducing glycerol into the ligand solution to get a final concentration of 30 %. The equilibration time in glycerol was 5 hours. The crystals were then frozen in liquid nitrogen. Diffraction data with these crystals was obtained at the Advanced Photon Source, Argonne National Laboratory (19-BM-D).

### Crystallographic data analysis

All the collected datasets were indexed, integrated, and scaled using HKL2000 using the programs Denzo and Scalepack^[19]^. The program Scalepack2MTZ was used to generate a reflection file in the CCP4 suite of programs ^[20, 21]^. Phasing information for both the structures was obtained using the phaserMR program on CCP4^[22]^. A high resolution native structure (PDB ID: 4I64) was used as a search model for the structure of the enzyme bound to mevaldehyde and Coenzyme A. For the structure obtained with the enzyme bound to mevalonate, Coenzyme A and NAD, a previously obtained quaternary complex structure was used as a search model without the ligands included. The search model was also solved via molecular replacement using a native pmHMGR structure. Structure files for the ligands were obtained using the HIC-UP server or using eLBOW from the Phenix suite of programs^[23-26]^. All parameter files have also been generated using eLBOW.

LigandFit from the Phenix was used to fit new ligand molecules in the generated 2Fo-Fc density^[24, 27]^. The structure was then run through a rigid body refinement using the program refmac5 in CCP4 followed by a simulated annealing (Cartesian) refinement using the Phenix.refine function ^[28]^. The structure was then run through several refinements using phenix.refine after using Molprobility and Real-Space Correlation to correct errors after each refinement until convergence of R_work_ and R_free_ values was observed^[29, 30]^. The program composite_omit_map in Phenix was used to generate omit maps for the ligands at the enzyme’s active site^[31]^.

## Results

### HMGR crystal structure soaked with mevalonate, CoA and NAD

Previous structural studies with dithio-HMG-CoA and NADH have shown that the enzyme is able to undergo major conformational changes to facilitate the enzymatic reaction and form reaction intermediates within the crystal^[9]^. In order to further understand the reaction steps 3 and 4 that lead to formation of mevaldehyde and mevaldyl-CoA formation, respectively (Fig. 1), we were interested in running the reaction within the crystal with mevalonate, CoA and NAD to observe the oxidative acylation of mevalonate.

**FIGURE 1.**
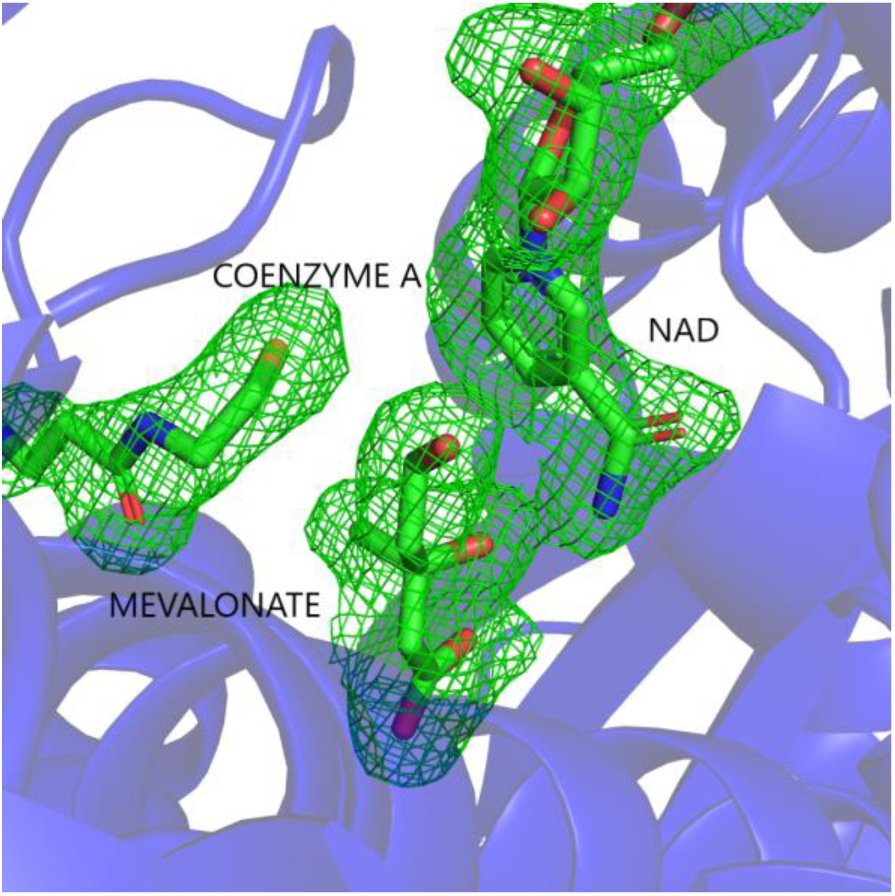
X-ray structure for HMGR crystals soaked with mevalonate, coenzyme A (CoA) and NAD highlighting the active site region with omit map density generated for the substrate and cofactors.

To investigate whether turnover can be observed in a mevalonate, CoA and NAD bound structure, we obtained a 2.2 Å structure soaked with the substrate (mevalonate) and cofactors (NAD and CoA). HMGR crystallizes as a homodimer in the asymmetric unit, with one catalytic site fully occupied and the other only bound to mevalonate due to the increased exposure to crystal contacts.

Electron density was observed for mevalonate, CoA and NAD but not for the thioester region between mevalonate and CoA that would denote the bond formation in the thiohemiacetal formation step indicating that no turnover has taken place.

### pH profile measurements in crystallization conditions

The lack of activity that was observed in the mevalonate, CoA and NAD bound crystal structure prompted us to determine the effect of the crystallization environment on the enzyme’s activity in this catalytic direction. In addition to the crystallization components, we also wanted to understand the effect of the crystallization pH (pH 6.7) on the activity observed in the crystallization environment. To determine the effect of the crystallization environment and pH on the enzymatic activity, the pH-activity profile for the rate of NADH production was measured in the crystallization buffer at saturated substrate concentrations. For comparison, we collected data using the enzymatic assay buffer used to characterize the pH-activity profile for the enzyme^[10]^.

The turnover of the enzyme was measured at various pH values from pH 4-12 and it was observed that the enzyme showed a sharp decrease in activity in the crystallization buffer in comparison to the enzymatic assay buffer (Fig 2.). The pH with maximum activity was also observed to shift from pH 9 to pH 10.

**FIGURE 2.**
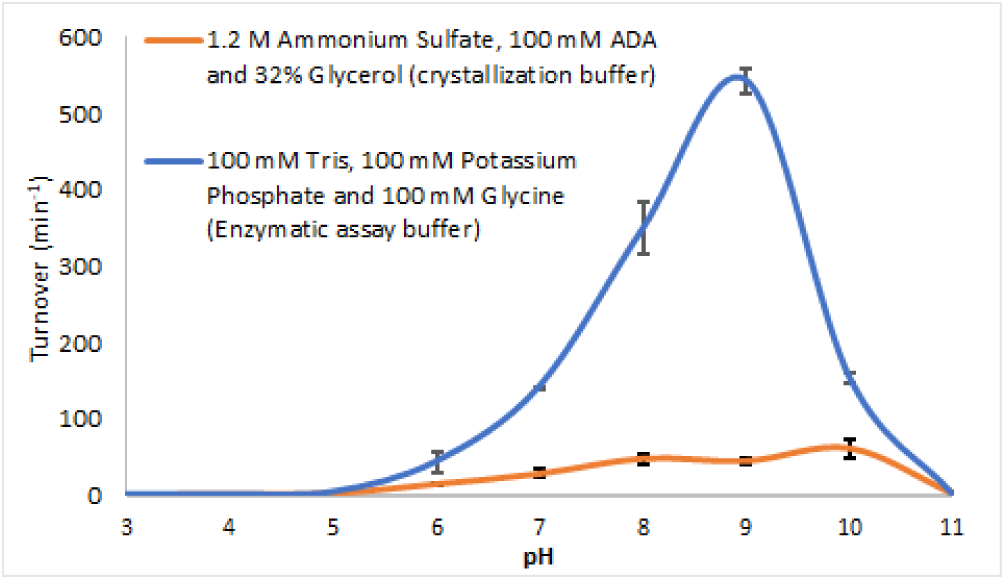
pH-activity profile showing the trendline for turnover of HMG-CoA reductase in enzymatic assay (Tris, Potassium Phosphate and Glycine) buffer (blue) and crystallization (Ammonium Sulphate, ADA and glycerol) buffer (orange) conditions.

### pH-activity profile in the presence of an alternate ammonium salt

The inhibitive effect from a high concentration of ions in the crystal has been previously observed in structural studies with Ribonuclease A and Ribonuclease 5^[32, 33]^. The inhibition in the Ribonuclease enzymes was alluded to be due to the binding of SO_4_^2-^ ions from the common crystallization salt Ammonium Sulfate.

To determine if the inhibitive effect observed with pmHMGR in the crystallization buffer is due to the presence of NH_4_^+^ or SO_4_^2-^ ions from Ammonium Sulfate, and to identify which of them produces an inhibitive effect, we measured the pH-activity profile of HMGR in the absence of sulfate with an alternate non-sulfate ammonium salt, ammonium chloride with the other crystallization buffer constituents.

We also observed an inhibition in the Ammonium Chloride, ADA and Glycerol environment in comparison to the enzymatic assay buffer (Fig 3.), although the inhibition is not found to be as strong as that observed in the crystallization environment. The enzyme also shows maximum activity at pH 10, like the crystallization buffer.

**FIGURE 3.**
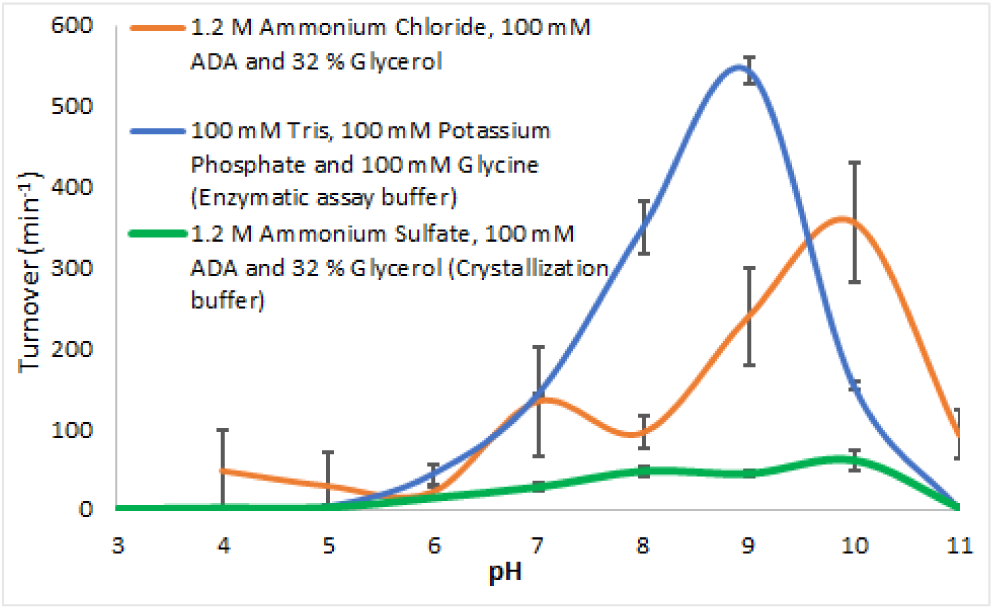
pH-activity profile showing the trendline for turnover of HMG-CoA reductase in an ammonium chloride, ADA and glycerol buffer (orange), crystallization buffer (green) and enzymatic assay buffer (blue).

Additionally, a combination of Ammonium Acetate, ADA and PEG-400 was also explored as an alternative ammonium-based non-sulfate buffer which differs from the crystallization buffer in the salt and cryoprotectant used. If exchanged with the HMGR crystal environment, the presence of PEG-400 has been proposed to prevent the lysis of the thioester bond which has been observed in structures of pmHMGR with HMG-CoA (PDB ID: 1R31), although the reasons for this are unclear^[9]^. A decrease in activity was observed in the ammonium acetate, ADA and PEG-400 buffer. The pH profile of the enzyme in this environment showed no activity until pH 9 and a sharp increase in activity after pH 9 (Fig 4.). A common trend of an increase in activity at a pH close to the pKa of Ammonium ions (9.3) is observed in the crystallization and non-sulfate ammonium salt buffers. The de-ionization of NH_4_^+^ ions is expected to occur with increasing pH. Hence, the increased activity at higher pH in the ammonium salt buffers points towards a potential inhibitive role of ammonium in the crystallization environment.

**FIGURE 4.**
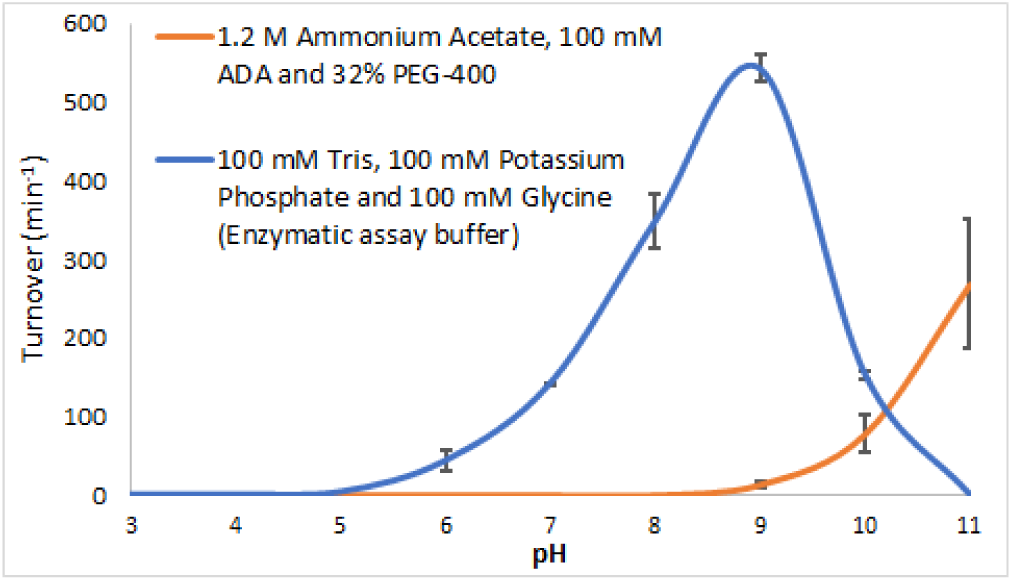
pH-activity profile showing trendline for turnover of HMG-CoA reductase in enzymatic assay (Tris, Potassium Phosphate and Glycine) buffer (blue) and Ammonium acetate, ADA and PEG-400 buffer (orange).

### pH-activity profile in the presence of non-ammonium salts

To further test if the inhibition observed in the crystallization environment is also observed in the absence of Ammonium, we measured the pH-activity profile of pmHMGR in the presence of a non-ammonium sulfate salt, Potassium Sulfate (Fig 5.). The buffer used also contained the other crystallization buffer constituents, ADA and Glycerol.

**FIGURE 5.**
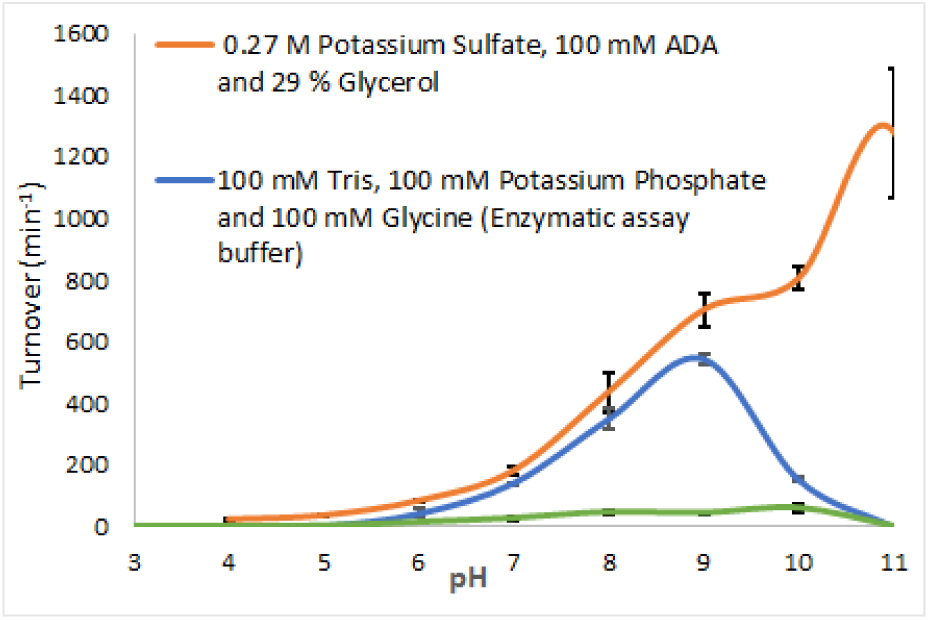
pH-activity profile showing trendline for turnover of HMG-CoA reductase in Potassium Sulfate, ADA and glycerol buffer (orange) in comparison to the enzymatic assay buffer (blue) and crystallization buffer (green).

In contrast to the ammonium salts, the potassium sulfate buffer did not show any reduction in activity when compared to the enzymatic assay buffer indicating that it is the ammonium and not sulfate present in the crystallization buffer that is reducing the activity of the enzyme.

### Changes in K_m_ for the enzyme ligands in the crystallization buffer

To further determine if the cause of inhibition in the crystallization buffer is a reduction in K_cat_ or if the crystal environment is also affecting the K_m_ values for the substrate (mevalonate) and cofactors (CoA and NAD) that produce HMG-CoA, we measured the K_m_ values for all the ligands in the crystallization buffer (Table 1).

**TABLE 1:** Km measurements for Coenzyme A, NAD and Mevalonate in Enzymatic assay conditions (0.1 M Tris-HCl, 0.1 M KCl, pH 8.1) and crystallization buffer.

| Buffer | CoASH (mM) | NAD (mM) | Mevalonate (mM) |
| --- | --- | --- | --- |
| Enzymatic assay buffer<br>(0.1 M Tris-HCl, 0.1 M KCl pH 8.1) | $0.05 \pm 0.002$ | $0.37 \pm 0.02$ | $0.22 \pm 0.01$ |
| Crystallization buffer (1.2 M Ammonium Sulfate, 100 mM ADA, 32 % Glycerol) | $0.03 \pm 0.004$ | $0.04 \pm 0.01$ | $0.06 \pm 0.01$ |
| $K_{m \text{ enz.}} : K_{m \text{ crys.}}$ | 1.67 | 9.25 | 3.67 |

The Km values observed indicate a significantly larger decline in the Km for NAD and mevalonate in the crystallization buffer in comparison to their previously characterized values[34].

The significantly larger reduction in Km specifically for NAD and mevalonate that are involved in the first hydride transfer step with the conversion of mevalonate into mevaldehyde (Fig 1) indicate the possibility of that this intermediate step being inhibited in the crystallization environment.

Since the assay being used to measure enzymatic activity measures the total NADH production from two hydride transfer steps (Scheme 3), the K_m_ values cannot be correlated directly with a single proposed reaction step. Hence, additional methods are required to investigate the reaction step we believe to be affected by the crystallization buffer in our kinetic studies.

**Scheme 3.**
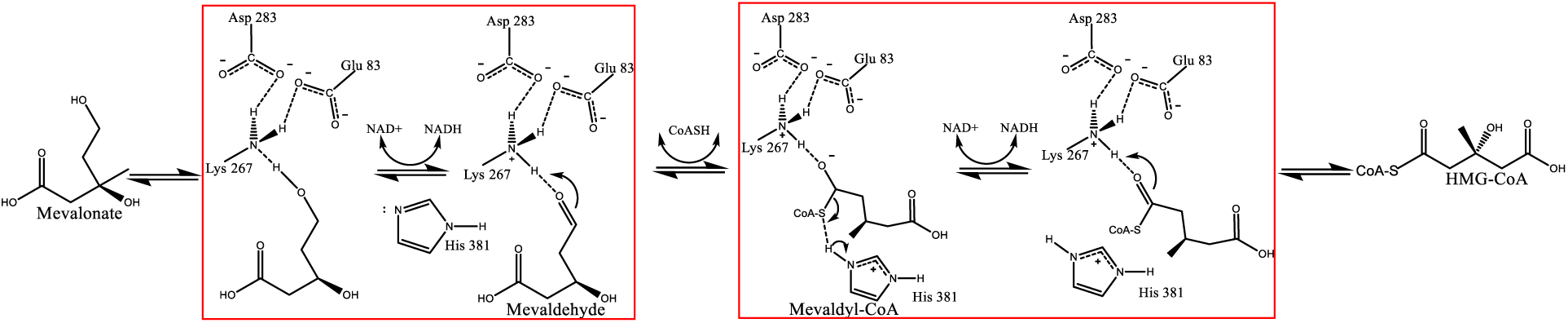
Schematic of the HMG-CoA reductase catalytic reaction for the conversion of mevalonate and Coenzyme A into HMG-CoA. Steps that involve a reduction of the NAD cofactor and are observed in the specific activity assay are highlighted in the red box.

### Turnover observed in crystals soaked with mevaldehyde and Coenzyme A

To further investigate the effect of the crystal environment on the hydride transfer step between mevalonate and NAD in pmHMGR, we crystallized the enzyme with the proposed post-turnover intermediate mevaldehyde and the cofactor CoA. The rationale behind looking into this ligand-soaked structure is to investigate if the enzyme is able to undergo turnover in the presence of mevaldehyde and CoA which would indicate that the crystal environment is inhibiting the reaction between mevalonate and NAD. Refined F_o_-F_c_ omit map density in the crystal structures show the emergence of an electron density in the C-S bond region between mevaldehyde and CoA with a bond distance of 1.85 Å indicating the formation of a thiohemiacetal intermediate (Fig 6A., Table 2). The C-S distance observed with the thiohemiacetal bond is found to closely match that of typical carbon-sulfur bonds (1.83 Å) ^[35]^. In comparing the observed thiohemiacetal conformation in this structure with the dithiohemiacetal structure observed in HMGR (*P. mevalonii*) upon soaking the slow substrate dithio-HMG-CoA and NADH (PDB: 4I4B), we see a close match in the overall configuration (Fig 6B.). The changes observed indicate that the enzyme can carry the reaction forward after the oxidation of mevalonate. The subsequent trapping of the thiohemiacetal is also observed in the crystal structure where the reaction intermediate does not undergo a second hydride transfer to form HMG-CoA. Since the same catalytic residues (Glu 83 and Lys 267) carry out both the hydride transfer steps with two separate NAD cofactors, it suggests that the second hydride transfer is also inhibited in the crystal via the observed NH_4_^+^ induced inhibition^[36]^.

**Table 2.**
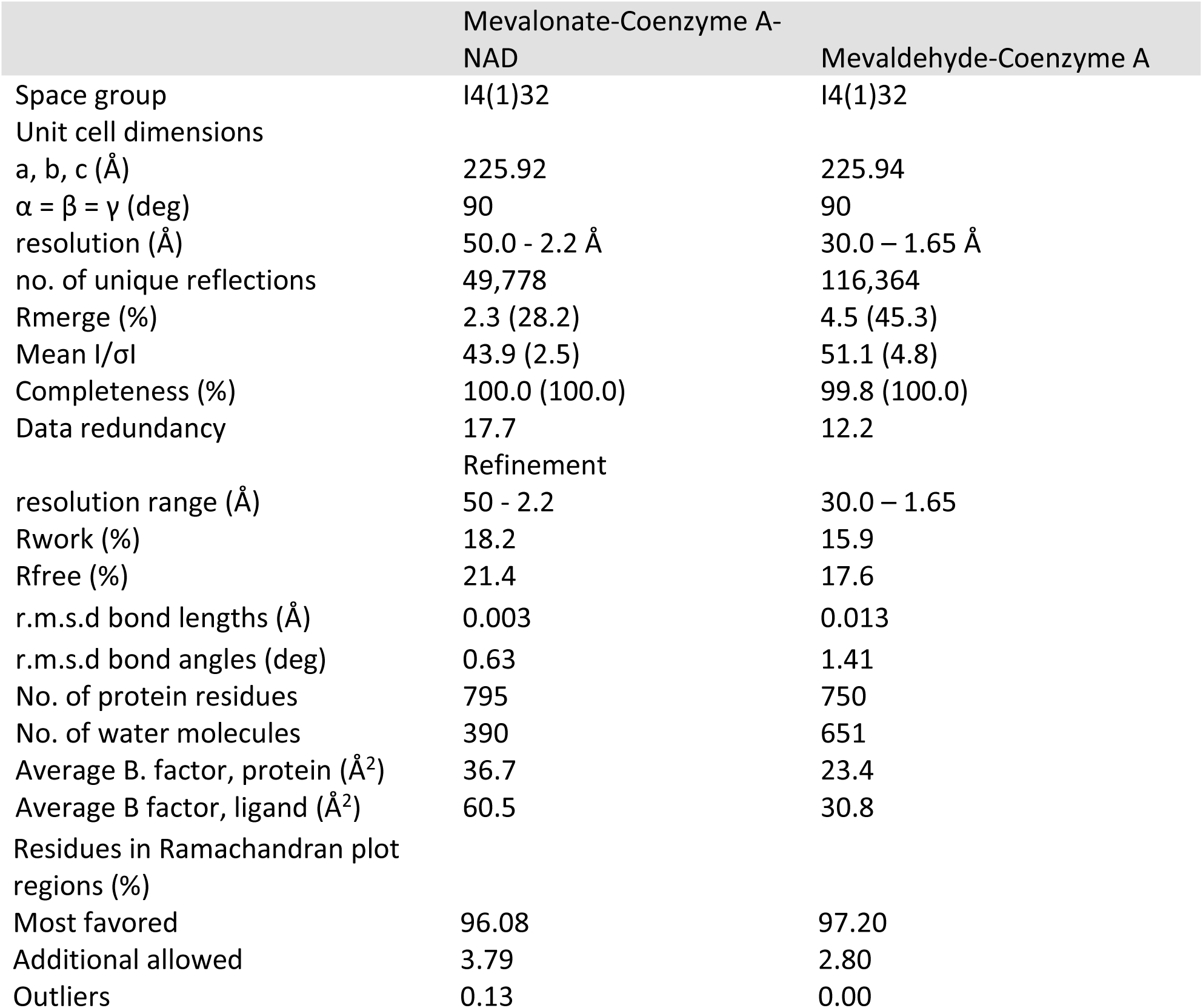
X-ray data collection and refinement statistics.

|  | Mevalonate-Coenzyme A-NAD | Mevaldehyde-Coenzyme A |
| --- | --- | --- |
| Space group | I4(1)32 | I4(1)32 |
| Unit cell dimensions |  |  |
| a, b, c (Å) | 225.92 | 225.94 |
| α = β = γ (deg) | 90 | 90 |
| resolution (Å) | 50.0 - 2.2 Å | 30.0 – 1.65 Å |
| no. of unique reflections | 49,778 | 116,364 |
| Rmerge (%) | 2.3 (28.2) | 4.5 (45.3) |
| Mean I/σI | 43.9 (2.5) | 51.1 (4.8) |
| Completeness (%) | 100.0 (100.0) | 99.8 (100.0) |
| Data redundancy | 17.7 | 12.2 |
|  | Refinement |  |
| resolution range (Å) | 50 - 2.2 | 30.0 – 1.65 |
| Rwork (%) | 18.2 | 15.9 |
| Rfree (%) | 21.4 | 17.6 |
| r.m.s.d bond lengths (Å) | 0.003 | 0.013 |
| r.m.s.d bond angles (deg) | 0.63 | 1.41 |
| No. of protein residues | 795 | 750 |
| No. of water molecules | 390 | 651 |
| Average B. factor, protein (Å<sup>2</sup>) | 36.7 | 23.4 |
| Average B factor, ligand (Å<sup>2</sup>) | 60.5 | 30.8 |
| Residues in Ramachandran plot regions (%) |  |  |
| Most favored | 96.08 | 97.20 |
| Additional allowed | 3.79 | 2.80 |
| Outliers | 0.13 | 0.00 |

**FIGURE 6:**
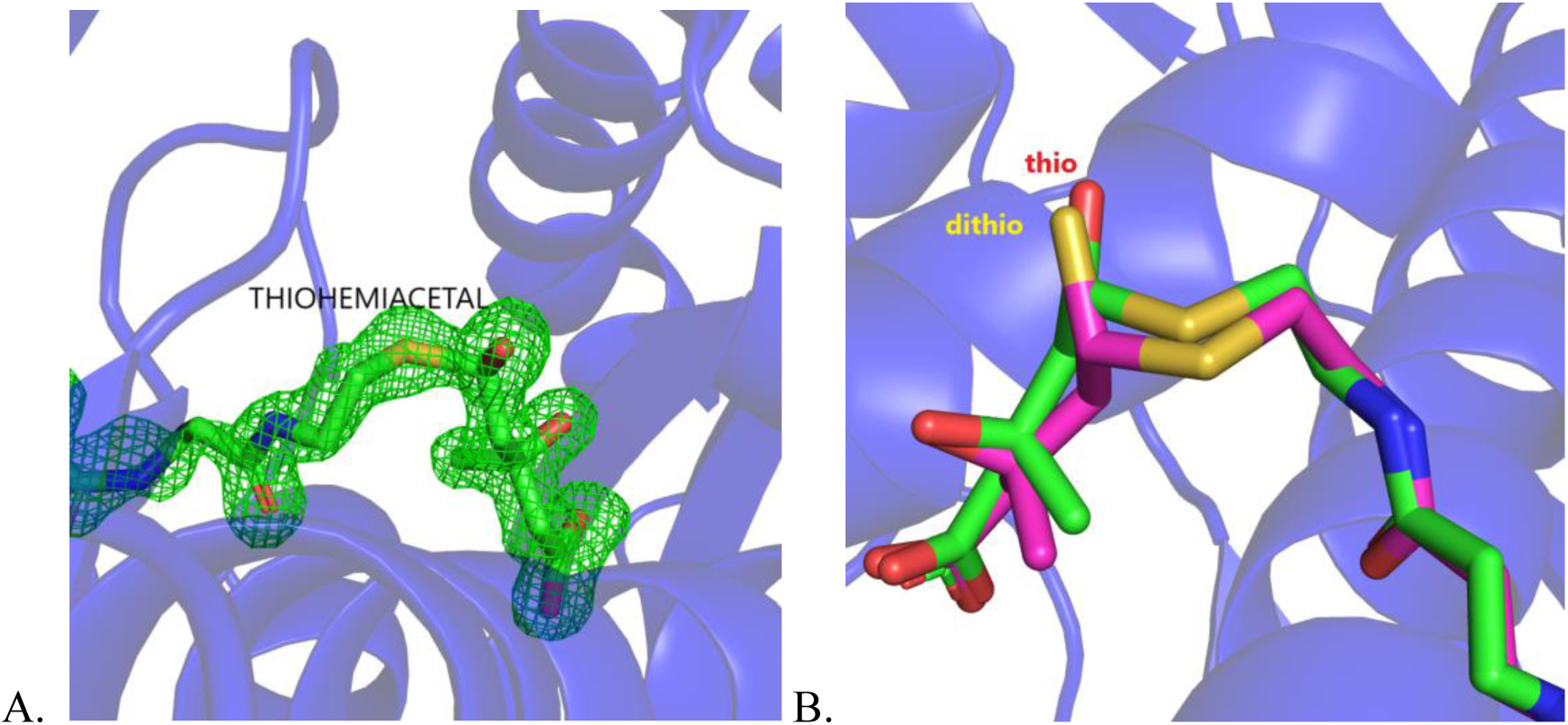
A. HMGR (*P. mevalonii*) active site showing bond density between mevaldehyde and Coenzyme A. 3σ F_o_-F_c_ difference electron density maps are shown in green. B. Thiohemiacetal conformation in structures with mevaldehyde and CoA (Green) in comparison with the dithiohemiacetal position obtained with dithio-HMG-CoA and NAD^+^ (Yellow).

## DISCUSSION

Numerous structural and biochemical studies have added to our current knowledge on the architecture of the HMGR active site and the residues that play a role in its formation and function. The structural studies have been performed using crystal structures of non-productive complexes, slow substrates, and selective ligands ^[6, 8]^. These studies have been able to elucidate important features such as the substrate and cofactor binding regions, conformational changes that allow catalysis to occur and some of the key catalytic residues involved in the enzymatic reaction. However, the conformational and chemical changes that allow the intermediates to form, the cofactor to be exchanged and the product to be released are still unclear. Further observations of these intermediary states of the enzyme along the reaction pathway are required to fully understand this complex enzymatic mechanism and confirm the proposed models for the enzymatic reaction.

Since the enzyme has been shown to undergo turnover within the crystal, we were interested in capturing the intermediary changes through crystal structures, particularly the conversion of mevalonate into mevaldehyde. Hence, we solved the structure of HMGR crystals with mevalonate, CoA and NAD with the intent of capturing post-turnover changes after the oxidative acylation of mevalonate. The quaternary complex only showed the pre-turnover state of the enzyme with no changes that would indicate the occurrence of a reaction with the crystal.

### Inhibitive role of Ammonium in the crystal environment

The inhibition of the enzymatic reaction that was observed in this crystal structure was further investigated by measuring the effect of the crystal constituents and pH on the enzyme. A pH-activity profile was measured in the crystallization buffer in which an inhibition of enzymatic activity was observed across pH 3-11. The observed trend in inhibition across pH in the crystallization buffer can be explained by the pKa of ammonium in ammonium sulfate solution.

The reported pKa for Ammonium ions in the Ammonium sulfate solution with the same ionic strength as that of the crystallization buffer is 8.74 – 8.77^[37]^. Presuming that ammonium ions act as an inhibitor, we would expect to see an increase in activity as the pH gets closer to and greater than the pKa of ammonium (pKa _NH4_^+^) due to its conversion to neutral ammonia. (Eq. 1) 

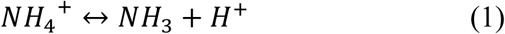

Additionally, we also expect the deionization of Ammonium to take place at a higher pH given that with increasing pH, we would observe an increase in OH^−^ ions which would promote and shift the equilibrium towards the formation of Ammonia (Shown in Eq. 2). 

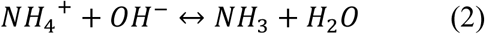

An overall increase in activity with increasing pH (pH 4-10) is observed in the crystallization buffer with the greatest activity at pH 10, where the pH > pKa _NH4_^+^ indicating that the observed inhibition is likely due to the presence of ammonium.

To investigate if the proposed inhibitive effect of NH_4_^+^ ions persists in the absence of the counter ion SO_4_^2-^ present in the crystallization buffer we measured the pH-activity profile of the enzyme in an alternate ammonium salt buffer consisting of Ammonium Chloride and the other crystallization buffer constituents, ADA and Glycerol. The enzyme continued to show a reduced turnover in comparison to the enzymatic assay buffer in the presence of Ammonium Chloride.

The increase in activity between pH 8-10 in the presence of Ammonium Chloride, ADA and Glycerol can also be explained through the dissociative properties of ammonium. At the concentrations tested, Ammonium in Ammonium Chloride is observed to have a pK_a_ value of 9.35-9.41^[38]^. At a pH greater than this pKa value, we expect a larger concentration of NH_4_^+^ to further dissociate and form NH_3_. The effect of de-ionization of ammonium is observed from pH 8-10, where we observe a reduction in inhibition as pH is closer to and greater than the observed pKa.

We also see an inhibition of the enzyme that correlates with the pKa of ammonium ions in another non-sulfate ammonium salt with Ammonium Acetate, ADA and PEG-400. Given that the pK_a_ of ammonium in ammonium acetate is 9.25, we expect the formation of NH_3_ to be greater in concentration after pH 9^[39]^. The expected increase in enzymatic activity is observed after pH 9 in Ammonium Acetate, ADA and PEG-400. This pH-activity profile like the ones observed with Ammonium Sulfate and Ammonium Chloride indicates that the observed inhibition is due to the presence of ammonium ions.

The same trend in activity is observed with increasing pH in all the ammonium salt buffers, which can be correlated with the changing ionization state of NH_4_^+^ ions in each buffer. This points to a pH-dependent inhibition from ammonium ions in the crystallization environment.

A variation in the degree of inhibition is observed in the presence of the different Ammonium salts. Possible reasons for the variation could be the difference in the pKa of the anions SO_4_^2-^, Cl^−^ and CH_3_COO^−^ that would lead to differences in the ionic interactions observed in each of the solutions, thereby affecting the interaction of NH_4_^+^ with the enzyme. Additionally, the different ions could have varying effects on protein stability as shown in the Hofmeister series where the different ions have been correlated with their effect on protein stability from least to most stable^[4]^: 

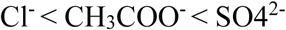

The effect of reduced activity with increased stability has been previously observed in T4-Lysozyme^[40]^. The enzyme pmHMGR undergoes drastic conformational changes upon ligand binding with the closure of the C-terminal flap domain at the ligand bound site^[8]^. Since these drastic conformations arenecessary for enzymatic activity in pmHMGR, it is possible that differences in the stability of the enzyme’s native state due to the ionic composition of the ammonium salt buffers tested contribute to the variation in the degree of inhibition that is observed.

To further investigate the effect of SO_4_^2-^ ions in the crystallization buffer, we measured the pH-activity profile of the enzyme in the absence of ammonium with potassium sulfate and the other crystallization buffer constituents (ADA and glycerol). No inhibition is observed within pH 4-11 in the presence of potassium sulfate in comparison to the enzymatic assay buffer.

The lack of inhibition in the buffers without Ammonium but with an alternate ion K^+^ also points to the inhibitive effect of ammonium ions in the crystal. It is important to note, however, that in the Potassium Sulfate buffer, the concentration of sulfate is 22 % of that observed in the crystallization buffer due to its lower solubility. However, significant inhibition continues to be observed in the presence of ammonium acetate and ammonium chloride where there are no sulfate ions present indicating that they do not reduce the activity in the crystal environment.

The lack of inhibition in the presence of Potassium Sulfate with ADA and Glycerol also points to the other crystallization buffer components (buffering agent and cryoprotectant) not having a role in reducing the activity in the crystallization environment.

Overall, the pH-activity profiles measured in various ammonium and non-ammonium salt buffers indicate that ammonium ions act as an inhibitor of the enzymatic reaction in the crystallization environment.

### K_m_ measurements in the crystallization buffer

The NH_4_^+^ dependent inhibition observed in the crystallization environment was further explored by measuring its effect on the ligand binding with the enzyme as reflected in kinetic parameters. To make this measurement, the K_m_ values of the substrate mevalonate and cofactors CoA and NAD that are involved in the production of HMG-CoA were measured. The significant reduction observed in the K_m_ values for mevalonate and NAD indicate that that the crystallization buffer could be inhibiting the step converting mevalonate into mevaldehyde since it involves the hydride transfer between NAD and mevalonate. The flattening of NADH production that is observed at lower ligand concentrations with NAD and mevalonate in comparison to the measurements obtained in the enzymatic assay buffer can be interpreted in conjunction with the observed evidence for the inhibition being dependent on ammonium binding in the crystal environment.

A proposed model that incorporates both results can be made considering the residues involved in the activity of HMG-CoA reductase. An examination of the active site suggests that the binding of NH_4_^+^ (Shown in Fig 7B.) to an anionic Glu 83 could inhibit the reaction step that requires a proton transfer from mevalonate to Glu 83 and a hydride transfer to NAD to produce mevaldehyde (Shown in Fig 7A.)^[8, 9, 11]^. The binding of NH_4_^+^ to Glu 83 sidechain could also be reducing the binding of mevalonate to the active site.

**Figure 7.**
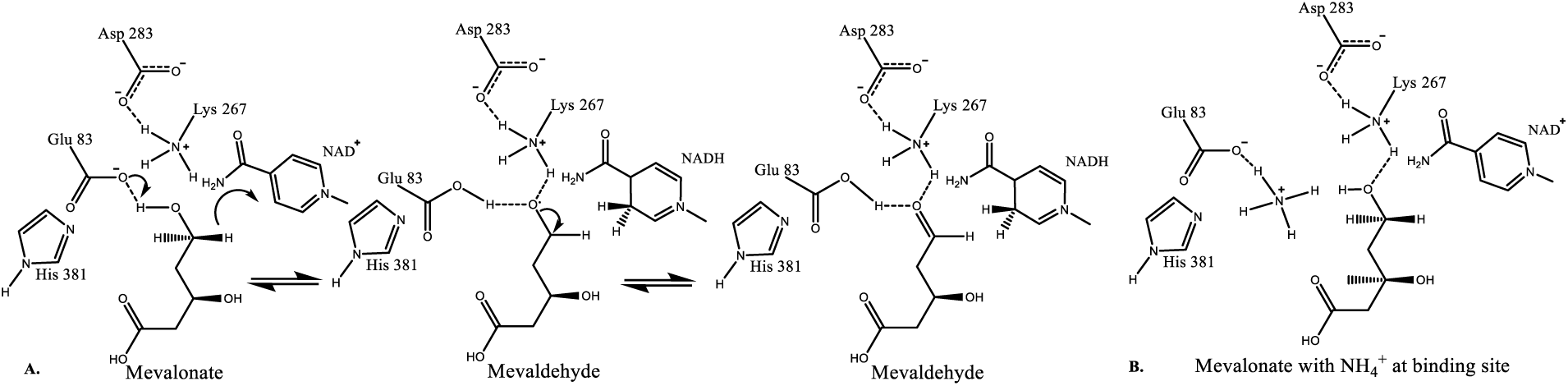
A. Schematic showing conversion of mevalonate into the proposed intermediate mevaldehyde. B. Model showing the proposed inhibition of Ammonium at the active site.

### Observance of turnover with mevaldehyde and Coenzyme A

The effect of inhibition on the first hydride transfer step was further shown with a comparison of the formation of the thiohemiacetal in the mevaldehyde and CoA bound structures to the initial substrate complex of mevalonate, CoA and NAD which shows no conversion to a hemiacetal. The observance of the thioester bond between mevaldehyde and CoA indicated that the enzyme can undergo turnover post hydride-transfer within the crystal indicating that the step being inhibited in the crystal is the oxidative acylation of mevalonate. The subsequent oxidation of the observed thiohemiacetal which also undergoes a hydride transfer at the catalytic site to form HMG-CoA also appears to be inhibited via the same NH_4_^+^ dependent mechanism resulting in an overall reduction in NADH production.

By identifying this pH-dependent inhibitive property of ammonium, we can now develop new methods and identify crystal environments to overcome this inhibition to obtain post-turnover complex structures of pmHMGR. These structures can then be useful in studying the structural and chemical changes that accommodate unexplored steps in the HMGR multistep reaction mechanism.

## ACKNOWLEDGEMENTS

We would like to thank the members of Hockmeyer Hall of Structural Biology for their support of this research.

